# Kcnb1-Kcng4 axis regulates Scospondin secretion and Reissner fiber development

**DOI:** 10.1101/2024.12.20.629661

**Authors:** Rasieh Amini, Ruchi P. Jain, Vladimir Korzh

## Abstract

The voltage-gated potassium channel Kv2.1 plays a role in the development of the ventricular system and the subcommissural organ in zebrafish. Here, a role for Kv2.1 in the secretion of the major component of Reissner’s fiber, Scospondin, was demonstrated. The results showed that Kv2.1 acts as a negative regulator of Scospondin secretion and Reissner fiber assembly. Kv2.1 regulates formation of Scospondin microfilaments and their assembly in Reissner fiber. Cholesterol playing a key role in Scospondin secretion. After the Reissner fiber is formed, it is detached from the hindbrain floor plate, where Scospondin produced initially. The tension of the fiber depends on its attachment to the subcommissural and flexural organs. In turn fiber tension affects the morphogenesis of these organs. This process of Reissner fiber formation depends on the input provided by the Hedgehog and Wnt/β-catenin signaling pathways on the anterior roof and floor plates.

## Introduction

Vertebrates can be roughly divided into two groups. Most vertebrates spend time in a horizontal position. They have the Reissner’s fiber (RF), an acellular structure that extends through almost the entire neural tube from the cerebral aqueduct through the fourth ventricle and spinal cord, terminating at the ampulla terminalis at the end of the central canal. In adults, the RF originates from the subcommissural organ (SCO), which develops as an enlargement of the anterior roof plate (RP) and ependyma near the entrance to the cerebral aqueduct (CA, Schoebitz et al., 1993;García-Lecea et al., 2017; Kiecker, 2018; Korzh and Kondrychyn, 2020; Nicholls, 1913; Rodríguez et al., 1998, 1992; Szathmari et al., 2013). RP develops under the influence of signaling in the dorsal neural tube, including BMP signaling, which has been associated with low cytoplasmic viscosity (Butler and Dodd, 2003; Yang et al., 2023) and Wnt/β-catenin signaling. One of several Wnts expressed by the RP, Wnt3 is evolutionarily conserved and associated with head development and cytoplasmic protrusion formation in Hydra and vertebrates (Holstein, 2024; Teh et al., 2015). The RF is under tension as it is anchored to the SCO, as well as to the caudal ends of the CA and the fourth ventricle (Woollam and Collins, 1980). The recent model suggested that starting from three days post fertilization (dpf) the RF in zebrafish larvae is a soft but tense elastic polymer (Bellegarda et al., 2023).

The giant matricellular protein Scospondin (Sspo) plays a central role in RF formation and is secreted by the SCO in adults and by both the SCO and the floor plate (FP) during development (Muñoz et al., 2019; Sepúlveda et al., 2021). The anterior FP forms the flexural organ (FO) at the caudal end of the CA (López-Avalos et al., 1997; Yang et al., 2021). Compared to the SCO, the development and function of the FO are much less understood. A question of evolutionary relationship to the infundibular organ of cephalochordates has finally been resolved in favor of the FO, suggesting its complex and ancient evolutionary history (Montiel and Aboitiz, 2018). Compared to FO, SCO is probably of later evolutionary origin. Sspo is conserved across species, as demonstrated by cross-species recognition by the anti-Sspo antibody, indicating its fundamental biological importance (Datki et al., 2023; Rodríguez et al., 1984; Yang et al., 2021). Several proteins produced by the SCO, FO, FP, and choroid plexus (ChP), have been implicated in RF formation [Clusterin (Clu), Galectin-1 (lgals1), and Chl1/Camel] (Hoyo-Becerra et al., 2005; Muñoz et al., 2019; Yang et al., 2021). Two of zebrafish *lgals1*-related genes (*lgals2a* and *lgals2b*) are expressed in ependyma (Thijssen et al., 2006). The Camel/Chl1a gain-of-function (GOF) induced ectopic RF formation, while its loss-of-function (LOF) hindered Sspo secretion and RF development (Yang et al., 2021). These observations emphasize the intricate molecular interactions involved in the RF’s formation and maintenance.

The development of the RF is initiated by the transient expression and secretion of Sspo by the FO, which subsequently extends along the anterior-posterior (A-P) axis in the FP. By 36 hours post-fertilization (hpf), Sspo expression ceases in the zebrafish FP. By 72 hpf, the RF acquires tension and detaches from the FP. The tense RF resonates while interacting with spinal sensory neuron as well as motile cilia that generate CSF flow (Bellegarda et al., 2023; Meiniel et al., 2008). The FP’s development relies on Nodal and Hedgehog (Hh) signaling originating from the embryonic shield and notochord (Le Douarin and Halpern, 2000; Lehmann and Naumann, 2005; Roelink et al., 1994; Sampath et al., 1998). The interplay of various genes within the Hh and Wnt signaling pathways is vital for regulating both the formation and function of the SCO and FP (Bach et al., 2003; Guiñazú et al., 2002; Lehmann and Naumann, 2005; Richter et al., 2001). Like the FO, which represents the most anterior FP, the SCO represents the most anterior RP, the dorsal signaling center that expresses multiple Wnts (Krauss et al., 1992; Mattes et al., 2012; Molven et al., 1991). This regulatory network underscores the complex molecular mechanisms involved in the development of RF-producing organs and highlights the importance of these pathways in embryonic development.

The activity of both the Wnt and Hh pathways depends on the presence of cholesterol, the versatile lipid and essential structural component of the plasma membrane (Nusse, 2003). It regulates membrane fluidity and is needed to form the lipid rafts involved in signal transduction (Pike, 2003). Cholesterol plays an important role in the Hh maturation and Wnt secretion (Beachy et al., 1997; Porter et al., 1996). It selectively activates the canonical Wnt signaling over non-canonical Wnt signaling (Sheng et al., 2014). The zebrafish Wnt3 transgenics were instrumental in demonstrating that the Wnt3 secretion is the cholesterol-dependent process (Ng et al., 2016). Using a combination of confocal microscopy and fluorescent correlation spectroscopy (FCS), the two main sources of Wnt3 in the developing brain were identified in the epithalamus and midbrain-hindbrain boundary (MHB) (Veerapathiran et al., 2020), which correlates with positions of the SCO and FO, correspondingly.

It has previously been shown that RF formation in the developing zebrafish occurs in two distinct phases. Initially, Sspo is expressed and secreted by the FP and later by the SCO. By 48 hpf, FP-associated *sspo* transcription is no longer detected except in the FO, and by 120 hpf it remains only in the SCO (Lehmann and Naumann, 2005; Meiniel et al., 2008). During development, zebrafish transgenics *sqet33mi2AEt* (hereafter referred to as ET33-mi2a) express the GFP in SCO, *etc.* This line carries the Tol2 transposon insertion in the *prom1a* intron and its GFP expression pattern recapitulates that of *prom1a* in the proliferative regions of the brain. (García-Lecea et al., 2017; Jászai et al., 2020; Jedrychowska et al., 2021; Jędrychowska et al., 2024; Kondrychyn et al., 2009; McGrail et al., 2010). Prom1/CD133 is a widely used stem cell marker involved in regulation of apicobasal cell polarity. It acts as the negative regulator of cholesterol metabolism (Lee et al., 2020) and is associated with the highly curved and prominent membrane structures such as cilia (Jászai et al., 2020; Pleskač et al., 2024). Several other developmental genes were linked to metabolism of cholesterol. *apoe* and *npc1* encode the cholesterol transporters. Mutations in these genes disturb cholesterol metabolism (Babin et al., 1997; Hu et al., 2022; Quelle-Regaldie et al., 2023).

An appropriately balance between the electrically active and modulatory subunits of the potassium voltage-gated channel Kv2.1 - Kcnb1 and Kcng4 is required for channel functions, including the maintenance of plasma membrane potential, formation of cholesterol-enriched lipid rafts, *etc*. (Deutsch et al., 2012; Johnson et al., 2019; Tamkun et al., 2007). Cholesterol is critical for secretion of ligands acting in developmental signaling (Abu-Siniyeh et al., 2016; Nusse, 2003). During development of the zebrafish BVS and inner ear, Kcnb1 and Kcng4b antagonize each other’s (Jedrychowska et al., 2021; Jędrychowska et al., 2024; Jędrychowska and Korzh, 2019; Shen et al., 2016). The loss-of-function (LOF) mutations of *kcnb1* and *kcng4b* cause BVS defects such as microcephaly and hydrocephalus. The hydrocephalic phenotype of the zebrafish Kcng4b LOF mutant is reminiscent of that of *hyh* mice deficient in α-Snap (Pérez-Fígares et al., 1998; Shen et al., 2016; Wagner et al., 2003). Like α-Snap, the Kcnb1 is involved in intracellular protein trafficking (Jędrychowska and Korzh, 2019; Jensen et al., 2017).

Since previous studies provided evidence that the defects of BVS in the *hyh* mice have been linked to abnormal development of the RF (Pérez-Fígares et al., 1998; Wagner et al., 2003), we decided to explore whether the Kv2.1 channel may play a role in the RF development.

## Results

### Kv2.1 regulates RF development

The combination of whole-mount anti-Sspo immunohistochemistry (IHC) and light-sheet confocal microscopy (LSCM) was used to study RF formation in *kcnb1* and *kcng4b* mutants on the background of ET33-mi2a transgenics expressing GFP in the SCO, FO, *etc*. (García-Lecea et al., 2017). When the 48 hpf wild-type controls (Fig. 1A;) were viewed in lateral view, the anterior RF (aRF) was detected between the SCO and the FO. The two parallel longitudinal signals were found posterior to the FO at the hindbrain level where the FP bends ventrally (Fig. 1A-C, white arrows; Fig. S1). This phenomenon has not been described before. The dorsal signal corresponds to the posterior RF (pRF), whereas the ventral signal is most likely located on the apical surface of the FP.

**Fig. 1.**
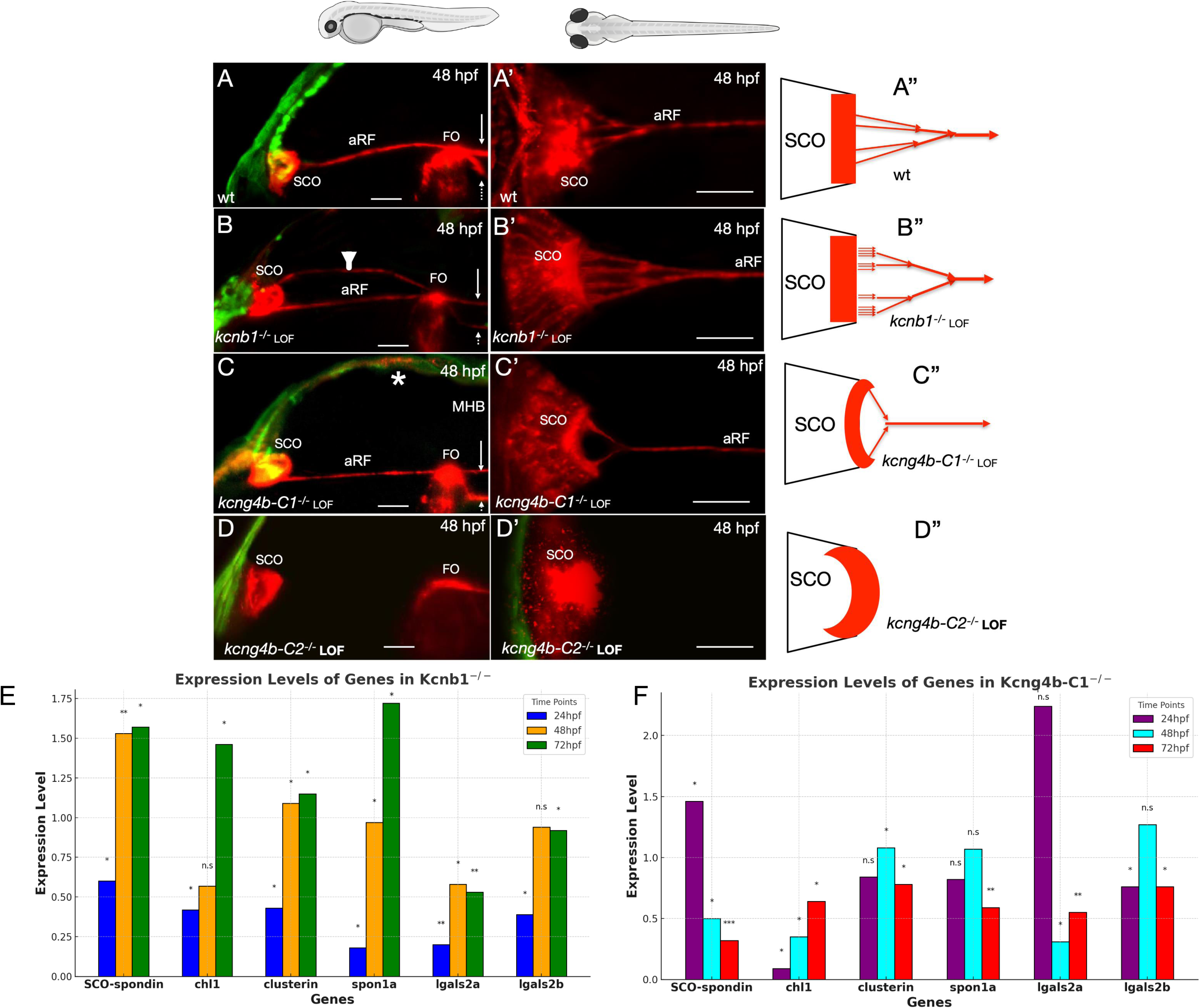
Mutations in genes encoding the Kv channel subunits Kcnb1 and Kcng4b affect the RF development. A, in the wild type 48 hpf embryo, the aRF connected the SCO and FO. A’ - several SCO-derived filaments merged to form the aRF. A” - scheme of A’. B - in the *kcnb1^-/-^* 48 hpf embryo, the aRF was duplicated. B’ - focus on the ventral aRF branch, where a number of SCO-derived filaments and nodes of merger increased compared to controls and an additional expression was present in the roof plate anterior to the midbrain-hindbrain boundary (MHB). B” - scheme of B’. C - in the *kcng4b-c1^-/-^* 48 hpf embryo, the aRF thickness was variable. C’ - a number of SCO-derived filaments was reduced to two with one node of merger. The apical SCO’ surface was concave. C” - scheme of C’. D - in the *kcng4b-c2^-/-^* 48 hpf embryo, the aRF was barely absent. D’ - no SCO-derived filaments were detected. The apical SCO’ surface was convex. D” - scheme of D’. Green - GFP expression in ET33-mi2a transgenics; red - anti-RF (AFRU) immunohistochemical staining detected by light-sheet confocal microscopy (LSCM). A-D - lateral view, A’-D’ - dorsal view, A”-D” - schemes based on A’-C’ images. Abbreviations: aRF - Reissner fibre, FO - flexural organ, pRF - posterior Reissner fibre. Scale bar - 20 um. E, F - developmental profiling of expression levels of the RF-associated genes detected by quantitative RT-PCR in the *kcnb1^-/-^* mutants (E) and *kcng4b-c1^-/-^* mutants (F).

In the 48 hpf *kcnb1^-/-^* mutant, the midbrain cavity representing the CA is reduced compared to the control (Shen et al., 2016). Here the two aRF branches were detected (Fig. 1B; arrowhead; Table 1). This could be due to an excess of Sspo secretion. This conclusion was supported by quantifying the expression of genes involved in RF development (Fig. 1D; Table 2). Of note, the *lgals2a* transcription was significantly lower compared to the control at 24 hpf, whereas *sspo* transcription after a moderate reduction at 24 hpf significantly increased at 48-72 hpf (Fig. 1D; Table 2).

**Table 1.** Mutations of genes encoding Kv2.1 subunits affect brain ventricular system and Reissner fiber.

| Experimental Group | Number of Samples Observed | Phenotype Description |
| --- | --- | --- |
| Wild Type (control, 48 hpf) | 20/20 | aRF detected between SCO and FO; two parallel longitudinal signals at hindbrain level (pRF and ventral FP apical signal) |
| *kcnb1*<sup>-/-</sup> mutant (48 hpf) | 6/20 | BVS reduced; two aRF branches detected |
| *kcng4b-cl*<sup>-/-</sup> mutant (48 hpf) | 20/20 | BVS increased; reduced aRF complexity |

**Table 2.** Mutations of genes encoding Kv2.1 subunits affect expression of genes implicated in RF formation.

|  | **Genes** | **Expression level ratio, *kcnb1*<sup>-/-</sup>/ wt, SD**<br>**24hpf** | **p value** | **Expression level ratio, *kcnb1*<sup>-/-</sup>/ wt, SD**<br>**48hpf** | **p value** | **Expression level ratio, *kcnb1*<sup>-/-</sup>/ wt, SD**<br>**72hpf** | **p value** |
| --- | --- | --- | --- | --- | --- | --- | --- |
| 1 | *SCO-spondin* | 0.6±0.08 | 0.05 | 1.53±0.36 | 0.01 | 1.57±0.6 | 0.05 |
| 2 | *chl1* | 0.42±0.35 | 0.05 | 0.57±0.18 | n.s | 1.46±0.04 | 0.05 |
| 3 | *clusterin* | 0.43±0.04 | 0.05 | 1.09±0.42 | 0.05 | 1.15±0.19 | 0.05 |
| 4 | *spon1a* | 0.18±0.09 | 0.05 | 0.97±0.16 | 0.05 | 1.72±0.09 | 0.05 |
| 5 | *lgals2a* | 0.2±0.16 | 0.01 | 0.58±0.19 | 0.05 | 0.53±0.38 | 0.01 |
| 6 | *lgals2b* | 0.39±0.23 | 0.05 | 0.94±0.03 | n.s | 0.92±0.04 | 0.05 |

| **N** | **Genes** | **Expression level ratio, *kcng4b-cl*<sup>-/-</sup>/ wt, SD**<br>**24hpf** | **p value** | **Expression level ratio, *kcng4b-cl*<sup>-/-</sup>/ wt, SD**<br>**48hpf** | **p value** | **Expression level ratio, *kcng4b-cl*<sup>-/-</sup>/ wt, SD**<br>**72hpf** | **p value** |
| --- | --- | --- | --- | --- | --- | --- | --- |
| 1 | *SCO-spondin* | 1.46±0.04 | 0.05 | 0.50±0.09 | 0.05 | 0.32±0.07 | 0.001 |
| 2 | *chl1* | 0.09±0.2 | 0.05 | 0.35±0.04 | 0.05 | 0.64±0.1 | 0.05 |
| 3 | *clusterin* | 0.84±0.19 | n.s | 1.08±0.04 | 0.05 | 0.78±0.26 | 0.05 |
| 4 | *spon1a* | 0.82±0.6 | n.s | 1.07±0.42 | n.s | 0.59±0.06 | 0.01 |
| 5 | *lgals2a* | 2.24±0.07 | 0.5 | 0.31±0.26 | 0.05 | 0.55±0.23 | 0.01 |
| 6 | *lgals2b* | 0.76±0.36 | 0.05 | 1.27±0.18 | n.s | 0.76±0.45 | 0.05 |

In the 48 hpf *kcng4b-c1^-/-^* mutant, which represents the hypomorphic Kcng4b loss-of-function allele, the BVS including CA is enlarged compared to the wild-type control (Jędrychowska et al., 2024; Shen et al., 2016). The aRF seems to be reduced (Fig. 1C, Table 1). This trend was emphasized in the dominant-negative *kcng4b-C2^-/-^* mutant, where the aRF was absent (Fig. 1D, D’, D”). These observations were supported by robust reduction of *sspo* transcription at 48 hpf compared to the control (Fig. 1E; Table 2). In contrast to *kcnb1^-/-^* mutant, the *lgals2a* transcription significantly increased at 24 hpf. Thus, the expression levels of *sspo* and *lgals2a* have changed in opposite ways in two mutants.

The dorsal view of the 48 hpf control embryo revealed a complex pattern of RF assembly. In the medial-lateral plane, the four individual microfilaments derived from the relatively straight apical SCO’ surface were detected. They merged at least twice before forming the single aRF (Fig. 1A’, A”). In the *kcnb1^-/-^* mutant, the four groups of 2-4 primary microfilaments derived from the apical SCO’ surface. These microfilaments formed the aRF following at least three rounds of merger (Fig. 1B’, B”). In contrast, in the *kcng4b-c1^-/-^* mutant the complexity of filamentous network was reduced. Here the aRF was formed by merging the two microfilaments derived from the apical SCO surface, which in contrast to controls was concave (Fig. 1C’, C”), whereas in complete absence of aRF this surface was convex (Fig. 1D, D’, D”).

The analysis of RF formation in the Kcnb1-Kcng4b mutants revealed that Kv2.1 activity regulates the RF development. It suggested that the detailed developmental study of zebrafish RF morphogenesis may reveal new information about the formation of the RF and related structures. Therefore, we used the same methodology to re-examine the early stages of the RF development in wild-type zebrafish.

### The two sources and two component parts of RF

The RF development takes place in parallel with significant morphological changes of the two sources of Sspo, the FP and SCO. The low level of *sspo* expression was detected by anti-Sspo IHC first at 16 hpf in association with anterior FP (not shown). At 18 hpf, the intensity of this signal increased (Fig. 2A). This signal likely represents the future FO (Olsson, 1956). At 22 hpf, the signal posterior to FO representing the posterior RF (pRF) became duplicated along D-V axis (not shown). At 30 hpf the duplicated signal extended along the hindbrain (Fig. 2B). The ventral signal which is clearly revealed by the anti-Sspo IHC (Fig. 2C, 3B) compared to Sspo-GFP *in vivo* (Fig. S1A) seems to align with apical surface of the hindbrain FP (Fig. S1B). The dorsal signal detected by both markers likely represents the pFP (Fig. 2C, S1A). At the midbrain level, only a single FO-associated signal was detected. From 30 hpf, the FO started to elongate along D-V axis and at 72 hpf acquired a hook-like shape with more intense staining dorsally (Fig. 2C, D; S1B).

**Fig. 2.**
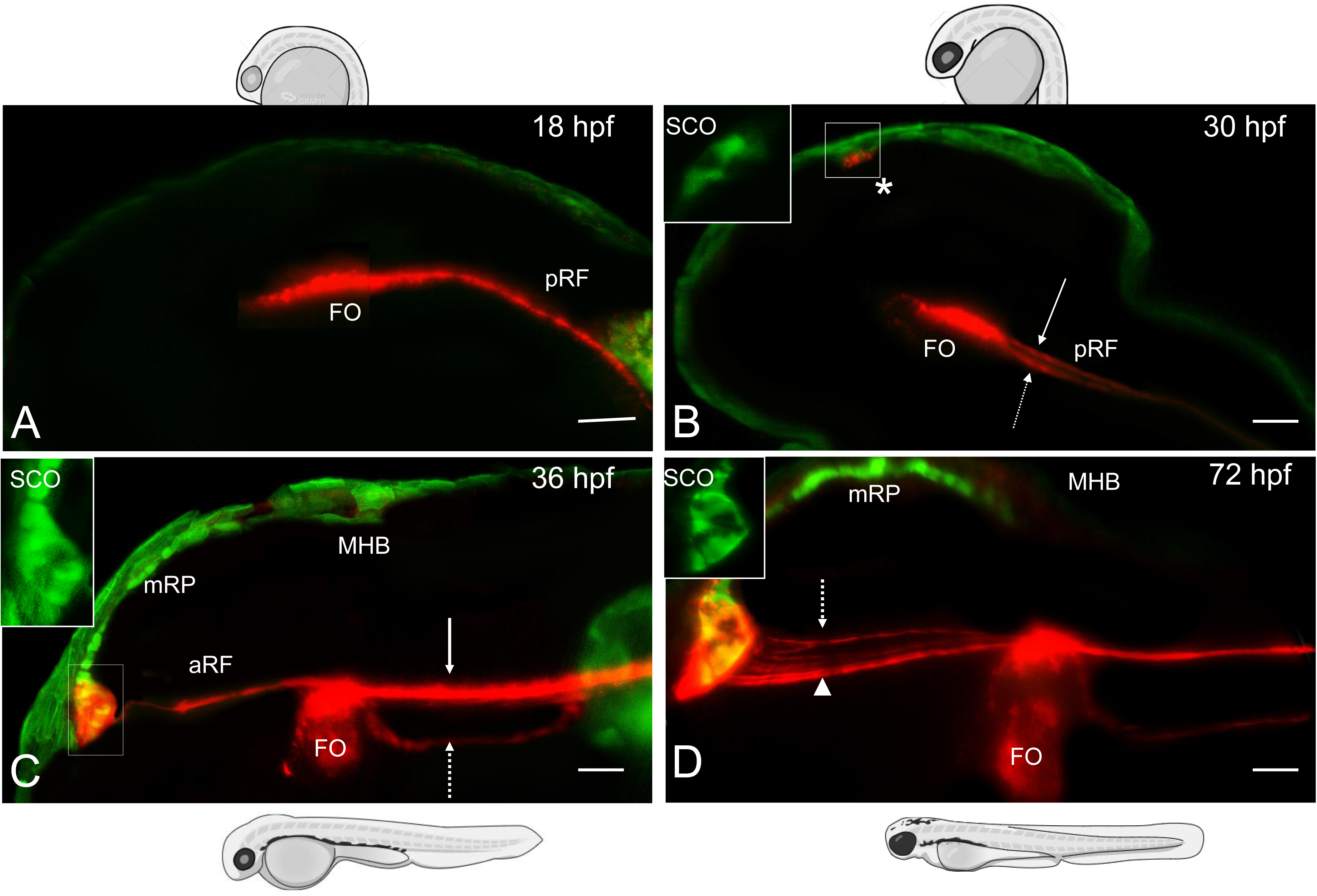
The steps in RF development. First, the pRF formed by the anterior FP. Second, the aRF formed by the SCO. Third, the pRF and aRF merged at the FO. A - in the 18 hpf embryo, the anterior FP started expressing Sspo (AFRU+, red). B, at 30 hpf, the SCO started expressing Sspo (asterisk). At the anterior hindbrain level, the signal has duplicated. C, at 36 hpf, the FO acquired its characteristic shape with a bulk of staining dorsally. The separation of two signals posterior of FO continued. The additional staining corresponding to the aRF was detected between the SCO and FO. Here the signal was more intense anterior of FO, whereas the region adjacent to SCO was weakly positive for AFRU staining. D, at 72 hpf the intensity of aRF staining increased, several SCO-derived branches of aRF formed, the SCO and FO became elongated. For 48 hpf wild type control, see Fig. 1A, A’. All images are in the lateral view. Abbreviations: aRF - Reissner fibre, FO - flexural organ, MHB - mid-hindbrain boundary, mRP - midbrain roof plate, pRF - posterior Reissner fibre. Scale bar - 20 um.

According to the anti-Sspo IHC, the SCO became Sspo-positive at 30 hpf (Fig. 2B). At 36 hpf, the anterior SCO-derived signal (aRF) connected the SCO and FO. aRF and posterior, FP-derived signal (pRF) merged at FO (Fig. 2C). The intensity of RF staining further increased at 48 hpf (Fig. 1A). At 72 hpf, several SCO-derived branches of RF were detected merging at different A-P levels between the SCO and FO (Fig. 2D).

During the 30-36 hpf period when the aRF was forming, the SCO cells began to rearrange and elongate. These events changed the organization of the anterior RP from a single-cell layer to a triangular pocket, with the central portion extending most posteriorly (Fig. 2B-E, inset). As the embryo developed, the SCO continued to elongate along the A-P axis. This previously described phenomenon has been attributed to the mechano-elastic deformation caused by RF pulling (García-Lecea et al., 2017) (Yang et al., 2021) Thus, these results not only supported the idea of consecutive formation of the RF by the FO and SCO (Lehmann and Naumann, 2005; Meiniel et al., 2008). They also revealed that soon after its formation at the hindbrain level, the RF lost contact with the FP, whereas maintaining its attachment to the SCO and FO. Unlike the hindbrain FP, the FO remained in contact with the aRF and pRF (Fig. 2C, 3). Thus, the FO acts as a connection hub for aRF and pRF and, in addition to SCO, as an attachment point for RF.

### RF morphogenesis depends on Wnt/β-catenin signaling

The Wnt3 promoter drives the expression and secretion of transgenic fluorescent markers and GFP-tagged Wnt3 in the anterior RP, i.e. SCO (Ng et al., 2016; Cathleen Teh et al., 2015; Veerapathiran et al., 2020). The two-color IHC analysis of Sspo expression in Wnt3 transgenics demonstrated the co-expression of Wnt3-GFP and Sspo in FO and SCO (Fig. 4A-C). The panembryonic RNAseq has shown that expression of several genes in the Wnt/β-catenin pathway was affected in the *kcng4b^-/-^* mutant (Jędrychowska et al., 2024; Shen et al., 2016). *rspo1* encodes the positive regulator of the Wnt/β-catenin pathway (Gore et al., 2011) expressed in the RP and SCO (Thisse et al., 2001). *foxa3* encodes the transcription factor expressed in the FP and FO (Odenthal and Nüsslein-Volhard, 1998). *rspo1* was transiently up-regulated in 24 hpf *kcnb1^-/-^* mutant, but not in the *kcng4b-c1^-/-^* mutant (Table 3). *foxa3* expression was not affected significantly in the *kcnb1^-/-^* mutant but increased in the *kcng4b-c1^-/-^* mutant along with *lrp6* which encodes Wnt3 coreceptor. The expression of the non-canonical *wnt4* did not change and that of *wls* involved in Wnt secretion remained flat in both mutants. These results suggested that the Wnt/β-catenin pathway likely plays a role during RF development although its mode of action in the RP and FP may differ.

**Table 3.** Mutations of genes encoding Kv2.1 subunits affect the midline signalling.

| N | Genes | Expression level ratio, *kcnb1*<sup>-/-</sup>/wt, SD<br>**24hpf** | p value | Expression level ratio, *kcnb1*<sup>-/-</sup>/wt, SD<br>**48hpf** | p value |
| --- | --- | --- | --- | --- | --- |
| 1 | *wnt4* | 1.015± 0.17 | 0.05 | 0.91±0.12 | n.s |
| 2 | *wls* | 1.125±0.17 | 0.05 | 0.94±0.08 | 0.01 |
| 3 | *rspo* | 1.48±0.47 | 0.05 | 0.67±0.26 | 0.05 |
| 4 | *shh* | 0.895±0.15 | n.s | 0.785±0.30 | 0.05 |
| 5 | *foxa* | 2.22±1.72 | 0.01 | 0.71±0.41 | 0.05 |
| 6 | *gli* | 0.93±0.09 | n.s | 0.8±0.28 | 0.01 |
| 7 | *lrp6* | 1.16±0.22 | 0.05 | 0.69±0.23 | n.s |
| 8 | *foxa3* | 1.06±0.09 | n.s | 0.62±0.23 | 0.05 |

| N | Genes | Expression level ratio, *kcng4b-CI*<sup>-/-</sup>/wt, SD, **24hpf** | p value | Expression level ratio, *kcng4b-CI*<sup>-/-</sup>/wt, SD, **48hpf** | p value |
| --- | --- | --- | --- | --- | --- |
| 1 | *wnt4* | 1.15±0.22 | 0.05 | 0.96±0.05 | 0.05 |
| 2 | *wls* | 1.05±0.07 | 0.05 | 0.97±0.04 | 0.05 |
| 3 | *rspo* | 0.945±0.07 | n.s | 0.67±0.34 | 0.05 |
| 4 | *shh* | 0.84±0.22 | n.s | 1.49±0.69 | 0.001 |
| 5 | *foxa* | 1.87±1.23 | 0.5 | 1.3±0.42 | 0.01 |
| 6 | *gli* | 1.87±1.23 | 0.01 | 1.56±0.79 | 0.001 |
| 7 | *lrp6* | 2.98±1.4 | 0.01 | 1.9±1.27 | 0.001 |
| 8 | *foxa3* | 1.38±0.54 | 0.05 | 3.51±3.55 | 0.001 |

It was shown that the tankyrase inhibitor (XAV939), which blocks the Wnt/β-catenin signaling, also inhibits the K^+^ current and cell proliferation *in vitro* (Huang et al., 2009; Ng et al., 2024). Given the co-expression of Wnt3 and Sspo in SCO, we asked whether RF would develop when the Wnt/β-catenin signaling is blocked? When the 24 hpf zebrafish embryos were treated with XAV939, the aRF and pRF were absent in 48 hpf embryos (Fig. 3D, E; Table 4). This phenotype seems to be more severe compared to that in the hypomorphic *kcng4b-c1^-/-^* mutant (Fig. 1C-C”). In contrast, after activation of the Wnt/β- catenin signaling by Wnt/β-catenin agonists [LiCl (Fig. 4F, G) or laduviglusib (CHIR-99021, Fig. 4H)], the aRF-associated signal expanded along the D-V axis. The SCO-derived network of Sspo+ microfilaments appears denser (Fig. 4G) compared to the control (Fig. 1D), which appears to be similar to the *kcnb1^-/-^*phenotype. Thus, the effect of Wnt/β- catenin activity on aRF inversely correlated with that of Kv2.1. In contrast to Kv2.1, the Wnt/β-catenin seems to act as a positive regulator of aRF.

**Fig. 3.**
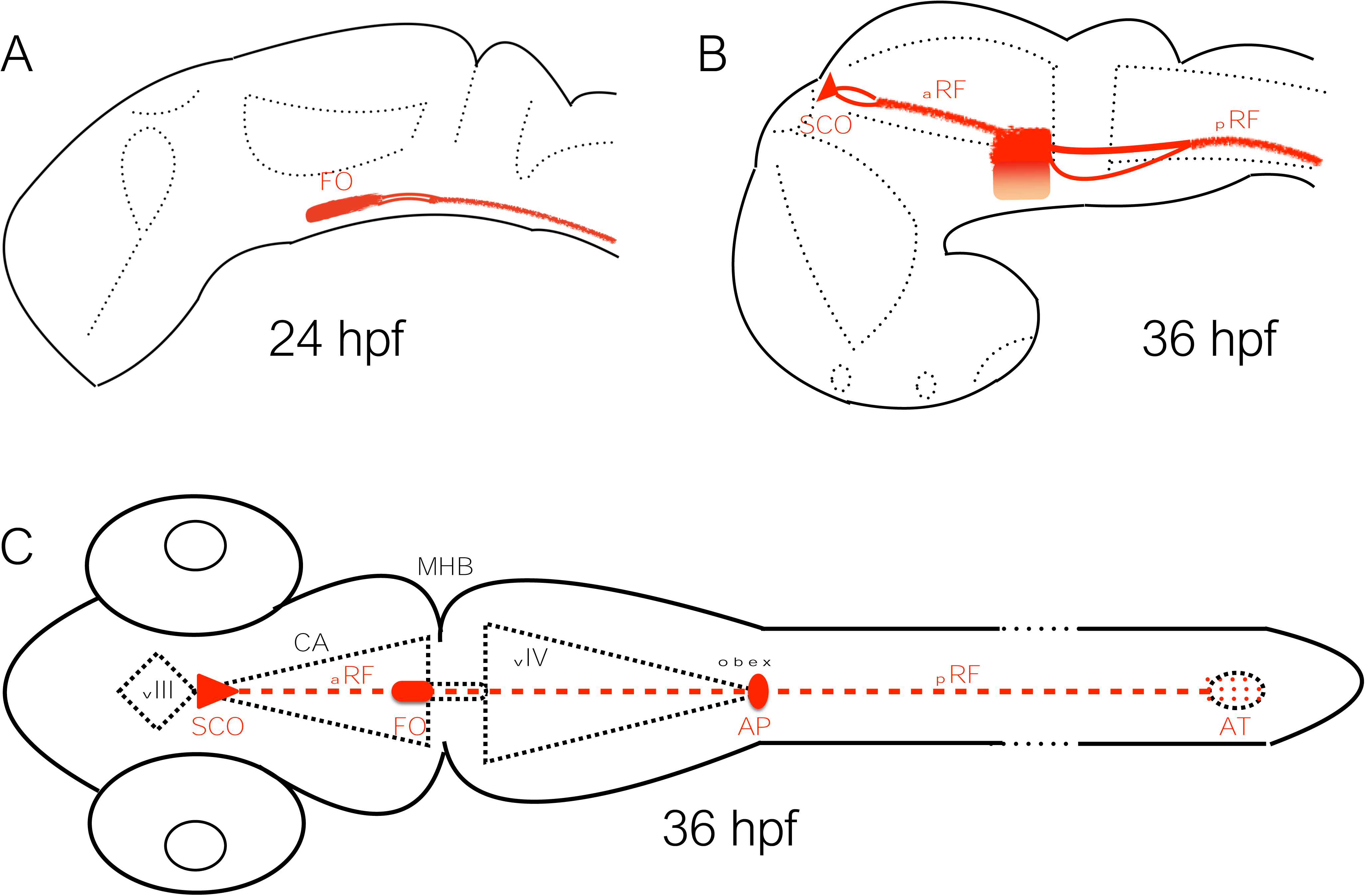
Schematic presentation of the CVOs (SCO, FO, AP) and RF. A, the FO/FP-derived RF (pRF) forms on an apical surface of FP. The AFRU+ signal is duplicated posterior to FO with the dorsal signal representing the pRF and the ventral one, the AFRU+ material on the apical surface of FP, which bends ventrally. B, the SCO-derived RF (aRF) forms in the cerebral aqueduct (CA). C, the RF is attached to the SCO and FO. It passes by the AP. These three CVOs are associated with the BVS narrows - CA and obex.

**Fig. 4.**
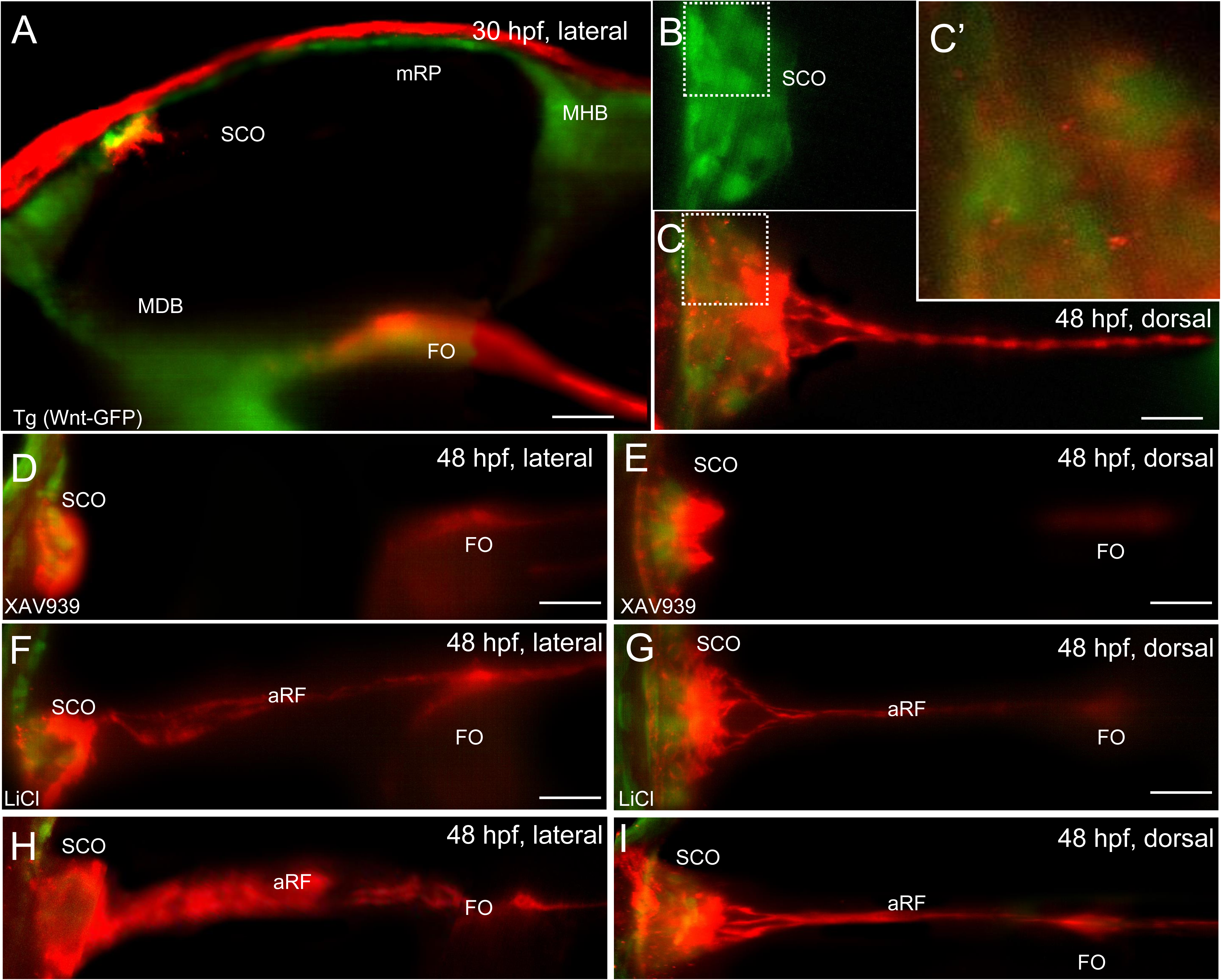
Wnt/β-catenin signaling regulates aRF formation. The SCO cells co-expressed Wnt3 and Sspo. A - the 30 hpf Wnt3-GFP transgenics (green) stained with the AFRU antibody (red). B - the SCO of the 48 hpf Wnt3-GFP embryo. C - the SCO of the same 48 hpf Wnt3-GFP transgenics stained with AFRU antibody, C’ - the 3x blowup of the boxed area (B, C). Wnt3-GFP and AFRU localize in the same cell, the GFP is mainly in perinucleal space and AFRU targets the peripheral cytoplasm; D, E, the pan-Wnt inhibitor (XAV939) blocks the development of the aRF and pRF both without noticeably affecting the SCO’ Sspo level. F, G, H - the Wnt/β-catenin agonists [LiCl (F, G) or laduviglusib (CHIR, H) increased the aRF- associated signal along the D-V axis. A, D, F, H - lateral view, B, C, E, G - dorsal view. Scale bar - 20 um.

**Table 4.** RF phenotype caused by experimental treatment.

| **Experiment** | **N embryos** | **N embryos without**<br>**aRF** | **N embryos with**<br>**scoliosis** |
| --- | --- | --- | --- |
| AFRU Injection | **90** | **20** | **15** |
| Kr15-16 Irradiation | **60** | **20** | **12** |
| Lrp5 morphants | **50** | **7** | **5** |
| Cyclopamine | **45** | **40** | **30** |
| XAV939 | **45** | **36** | **25** |

### RF morphogenesis depends on Hh signaling

Shh is the axial mesoderm-derived morphogen that acts downstream of the Nodal signaling to ventralize the neural tube and develop the FP (Krauss et al., 1993; Roelink et al., 1994; Sampath et al., 1998; Schier et al., 1997; Strähle et al., 2004). It has been shown previously that the Nodal deficiency affects the expression of Sspo in the FP, but not in the SCO (Lehmann and Naumann, 2005). Therefore, it is of interest to re-examine which aspects of RF formation depend on Hh activity. The Hh inhibitor cyclopamine was used to efficiently block the Hh activity and an agonist of Hh, purmorphamine (PMA), was used to increase the Hh signaling in the stage-specific manner during 24-48 hpf period (Winata et al., 2009).

Moderate cyclopamine-mediated inhibition of Hh signaling resulted in absence of aRF and reduction of Sspo staining at the FP and FO. The apical surface of the FO shrunk along the A-P axis and the duplicated signals in the hindbrain were reduced to such extent that the ventral signal was no longer detectable. In contrast, the intensity of SCO-associated signal increased (Fig. 5A-D). The PMA treatment did not cause any major changes compared to the wild-type control although the curvature of the hindbrain FP increased (Fig. 5E, H). Thus, the Hh signaling plays a critical role in the RF formation by regulating the FP- and SCO-related Sspo secretion.

**Fig. 5.**
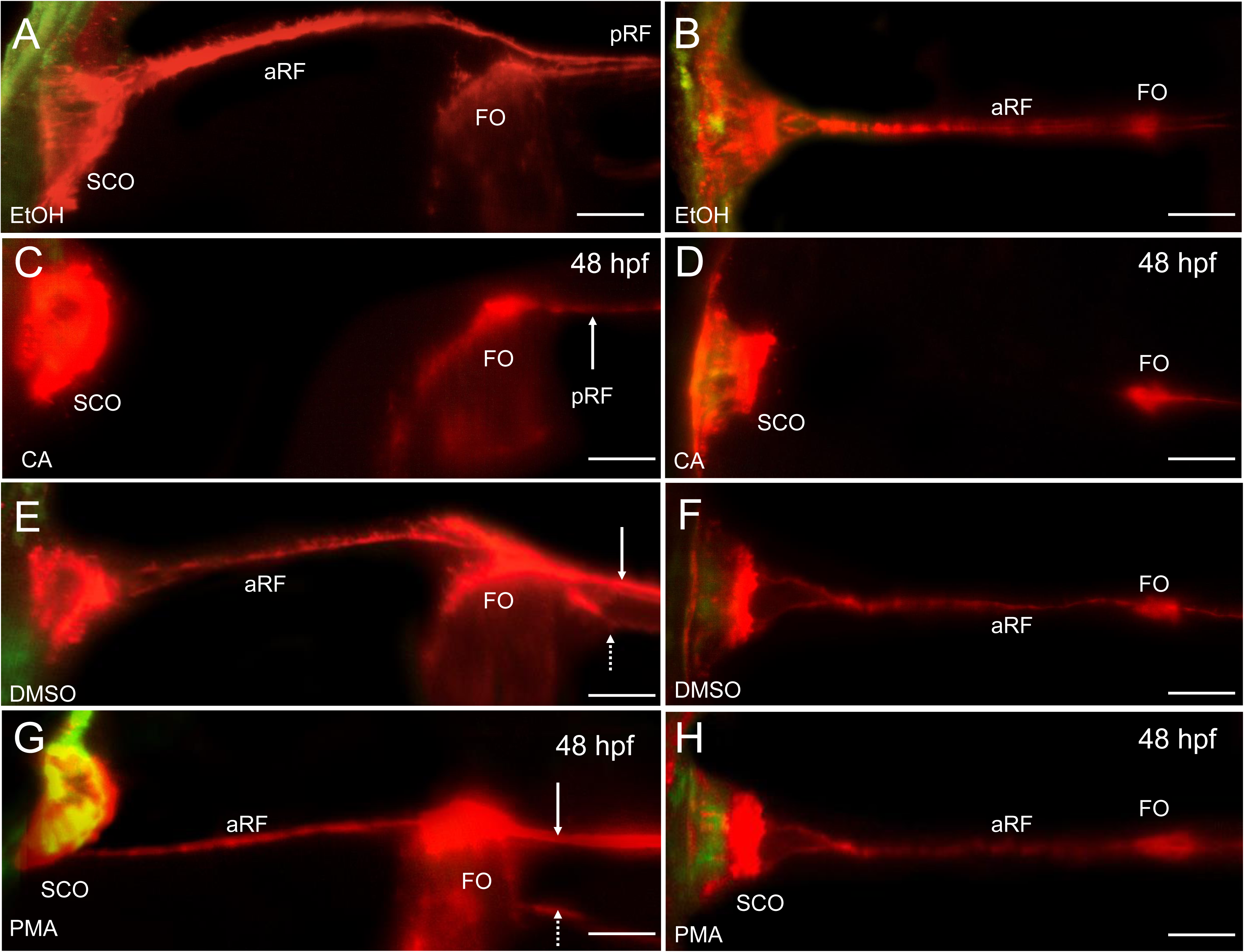
Hh signaling regulates RF formation. A, B, cyclopamine (CA) treatment (EtOH) blocks aRF development (48 hpf). The SCO ventrolateral surface is intensely labelled by AFRU ab. The apical FO surface shrinks and the signal at the FP apical surface is not detectable. A, lateral view, B, dorsal view. Of note is the inhibitory effect of EtOH on merger of the aRF and pRF at the FO (A) G, H, purmorphamine (PMA, DMSO) have no obvious effect on the RF development. The AFRU signal is concentrated closer to the SCO apical surface unlike that in CA-treated embryos. Of note is the inhibitory effect of DMSO of merger of microfilaments (F, H). Scale bar - 20 um.

### RF morphogenesis depends on cholesterol

The RNAseq data suggested that *prom1a* expression was affected in the zebrafish *kcng4b-tr* LOF mutants, which may have an impact on cholesterol metabolism (Jędrychowska et al., 2024). Given the role of canonical Wnt signaling in aRF formation, we asked a question whether cholesterol is required for the Sspo secretion and RF formation? We previously showed that the acute mβCD-mediated depletion of cholesterol in zebrafish embryos decreased the Wnt3 association with cholesterol-dependent domains in the plasma membrane (Ng et al., 2016). Acute mβCD treatment blocked aRF formation in wild-type embryos (Fig. 6A, B, D, E). Long-term exposure to statins inhibits cholesterol biosynthesis (Signore et al., 2016). In wild-type embryos exposed to atorvastatin, the Sspo was produced by SCO, but the aRF failed to form (Fig. 6A, C, D, F). Thus, both the chemical depletion of cholesterol and blockade of cholesterol synthesis affected the aRF formation.

**Fig. 6.**
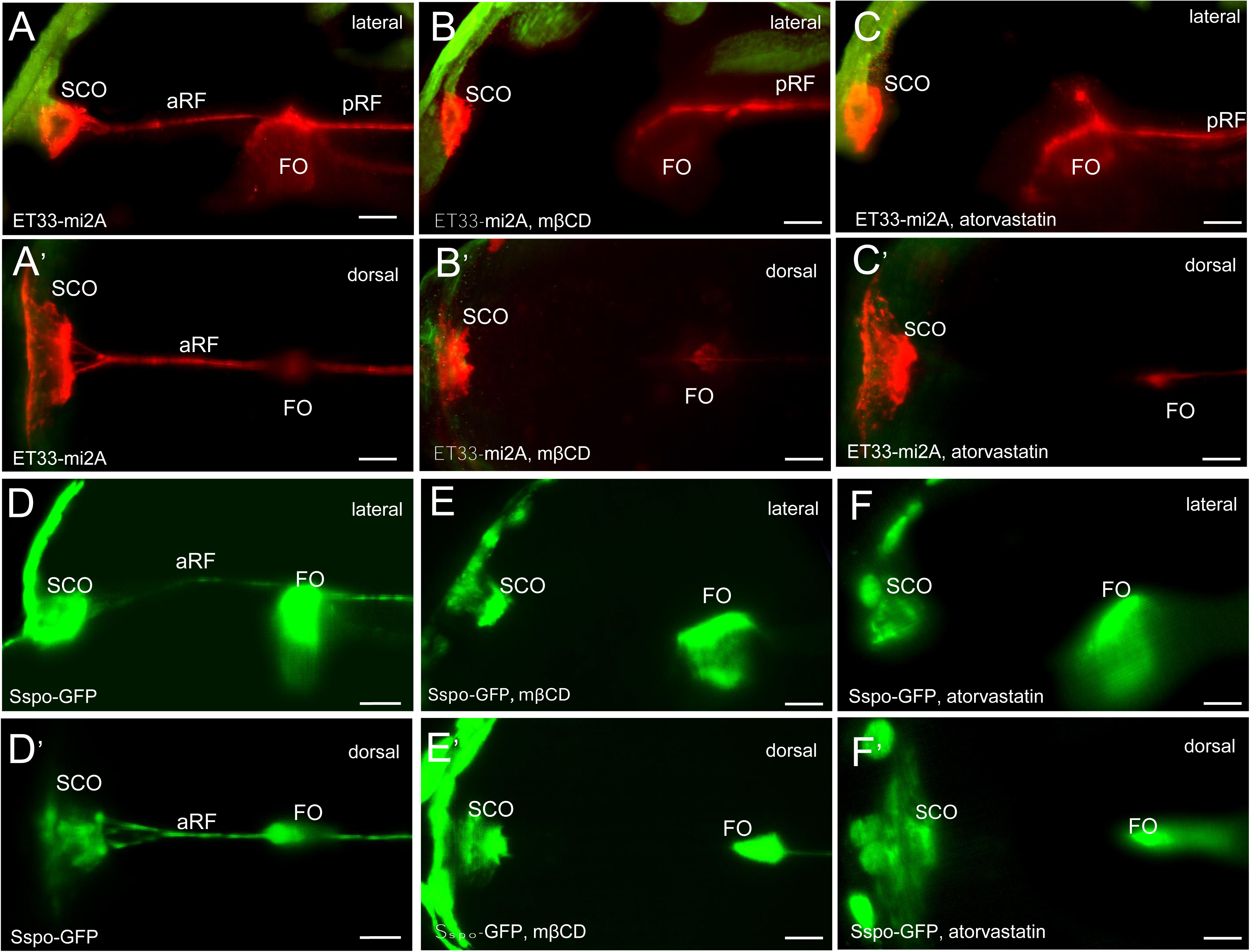
Cholesterol depletion affects the aRF formation. The aRF detected in fixed whole mount embryos by AFRU IHC (A-C) of in vivo in Sspo-GFP transgenics. A, A’, D, D’ - control; B, B’, E, E’ – 40 min treatment of 48 hpf embryos by 2.5 μM mβCD abolished the aRF in controls. C, C’, F, F’ - 24→48 hpf atorvastatin treatment (10 μM) abolished the aRF in controls. All images are in lateral view. Scale bar - 20 um.

Given the increased Sspo expression in *kcnb1^-/-^* mutant, the question was whether this excess was due to increased cholesterol and to what extent the Kcnb1 RF phenotype could be rescued by cholesterol reduction? Indeed, after the mβCD treatment of *kcnb1^-/-^*mutant the aRF was affected, but it was still detectable and maintained its orientation along the SCO-FO axis (Fig. 7A, C) unlike that after atorvastatin treatment which causes a more severe effect (Fig. 7B, D). Thus, in the cholesterol-depleted *kcnb1* mutants the reduced activity of Kv.2.1 rescued Sspo secretion and aRF formation, but chronic blockade of cholesterol synthesis by atorvastatin abolished this compensation.

**Fig. 7.**
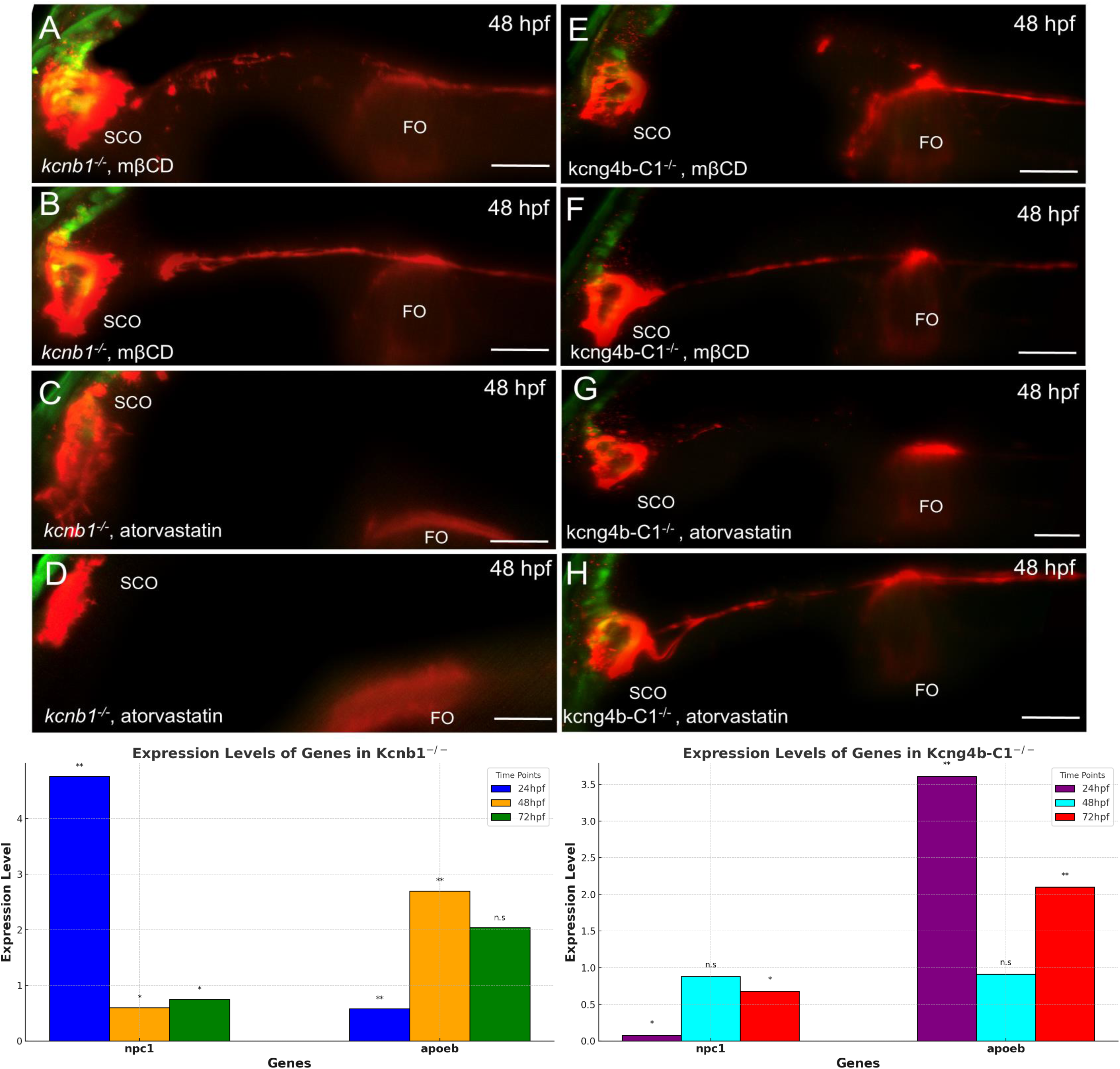
Cholesterol depletion affects the aRF formation in *kcnb1^-/-^* mutants. A-B – 40 min treatment of 48 hpf *kcnb1^-/-^* by 2.5 μM mβCD severed the RF organization. C, D - 24→48 hpf atorvastatin treatment (10 μM) abolished the aRF in *kcnb1^-/-^*. All images are in lateral view. Scale bar - 20 um. I, J - developmental profiling of expression levels of the cholesterol-associated genes detected by quantitative RT-PCR in the *kcnb1^-/-^* mutants (I), and *kcng4b-c1^-/-^* mutants (J).

In contrast, the weak phenotype of *kcng4b-c1^-/-^* mutant was not significantly affected by cholesterol depletion (Fig. 7E-H) as if the moderate increase in Kv2.1 activity prevented the complete suppression of Sspo secretion.

This suggested that cholesterol is required for Kv2.1-mediated regulation of Sspo secretion. To test this idea, the qRT-PCR was performed on *kcnb1* and *kcng4b* mutants to detect the transcription of genes known to regulate cholesterol metabolism. *npc1* encodes the cholesterol transporter that is deficient in the Neumann-Pick disease type 1 (Lin et al., 2018; Tseng et al., 2018). The expression of *npc1* was upregulated in *kcnb1* mutant and downregulated in *kcng4b-c1* mutant (Fig. 7I, J; Table 6) unlike the related *npc2* (Jędrychowska et al., 2024). *apoeb* encodes a protein involved in cholesterol efflux (Mahley, 1988; Monnot et al., 1999). The expression of *apoeb* expression was downregulated in the *kcnb1^-/-^* mutant (Fig. 7I), whereas in the *kcng4b-c1^-/-^* mutant it was upregulated (Fig. 7, J; Table 6).

**Table 5.** RF phenotype caused by cholesterol blockade.

| Experiment | N embryos | N embryos without aRF | N embryos with scoliosis |
| --- | --- | --- | --- |
| WT mβCD | 30 | 30 | 15 |
| mβCD (Tg Sspo-GFP/live imaging) | 30 | 30 | 21 |
| mβCD kcnb1-/- | 40 | 40 | 12 |
| mβCD kcng4b-C1-/- | 20 | 14 | 10 |
| WT Atorvastatin | 30 | 30 | 12 |
| Atorvastatin (Tg Sspo-GFP/live imaging) | 30 | 23 | 20 |
| Atorvastatin kcnb1-/- | 20 | 3 | 3 |
| Atorvastatin kcng4b-C1-/- | 20 | 8 | 6 |

**Table 6.** Mutations of genes encoding Kv2.1 subunits affect the expression of cholesterol transporters.

| N | Genes | Expression level ratio, *kcnb1*<sup>-/-</sup>/wt, SD<br>**24hpf** | p value | Expression level ratio, *kcnb1*<sup>-/-</sup>/wt, SD<br>**48hpf** | p value | Expression level ratio, *kcnb1*<sup>-/-</sup>/wt, SD<br>**72hpf** | p value |
| --- | --- | --- | --- | --- | --- | --- | --- |
| 1 | *npc1* | 4.76±0.15 | 0.01 | 0.60±0.02 | 0.05 | 0.75±0.04 | 0.05 |
| 2 | *apoeb* | 0.58±0.04 | 0.01 | 2.7±0.46 | 0.01 | 2.04±1.11 | n.s |

| N | Genes | Expression level ratio, *kcng4b-CI*<sup>-/-</sup>/wt, SD, **24hpf** | p value | Expression level ratio, *kcng4b-CI*<sup>-/-</sup>/wt, SD, **48hpf** | p value | Expression level ratio, *kcng4b-CI*<sup>-/-</sup>/wt, SD, **72hpf** | p value |
| --- | --- | --- | --- | --- | --- | --- | --- |
| 1 | *npc1* | 0.08±0.16 | 0.05 | 0.88±0.21 | n.s | 0.68±0.47 | 0.05 |
| 2 | *apoeb* | 3.61±0.70 | 0.01 | 0.91±1.03 | n.s | 2.10±0.76 | 0.01 |

The developmental analysis of Kcnb1-Kcng4b mutants revealed that the Kv2.1 activity regulates the RF morphogenesis, a complex and highly orchestrated process involving multiple signaling pathways including Wnt/β-catenin and Shh. RF formation is dependent on cholesterol metabolism. The Kv2.1 activity is determined by the antagonistic effects of its subunits, whose deficiency affects the expression of genes associated with RF development, cholesterol metabolism and Sspo secretion. Thus, Kv2.1 acts as a factor limiting Sspo secretion (Fig. 8).

**Fig. 8.**
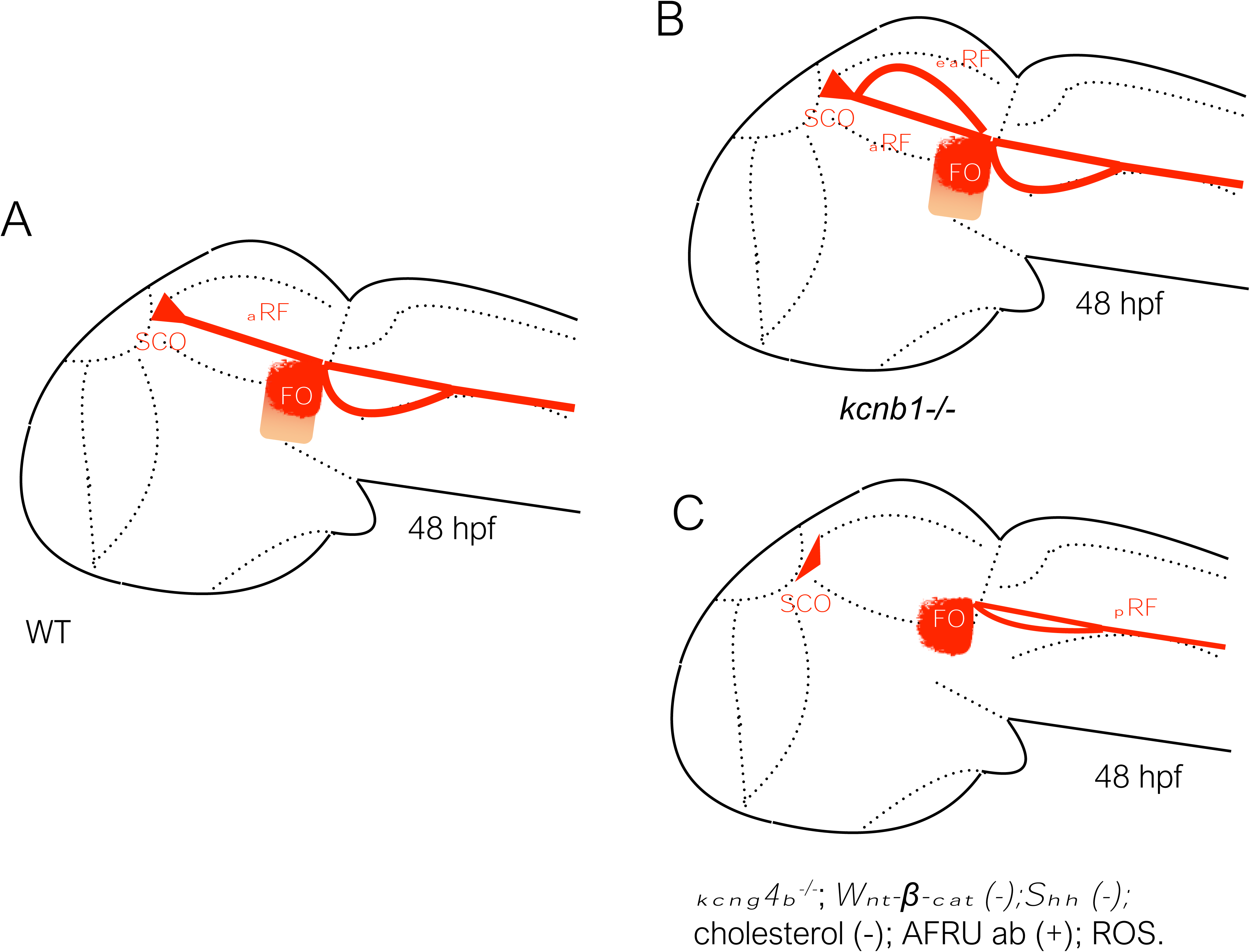
Multiple inputs regulate aRF morphogenesis. These are the signaling pathways (Shh, Wnt-β-cat) active in the FP and RP. Kv2.1 activity and cholesterol modulate Sspo secretion. The KillerRed-mediated production of ROS and microinjection of AFRU antibody block aRF formation with low efficiency.

## Discussion

Given the role of Kv2.1 in the development of BVS, where the channel subunits Kcnb1 and Kcng4b antagonize each other’s activity (Shen et al., 2016), the question has been raised as to how the changes in Kv2.1 activity may affect the RF? Previously, the BVS defects such as hydrocephalus in mammals (Cifuentes et al., 1994; Grondona, 1998; Lang et al., 2006; Pérez-Fígares et al., 1998; Rodríguez et al., 1990; Vio et al., 2008) and scoliosis in zebrafish (Grimes et al., 2016; Troutwine et al., 2020) have been associated with the RF abnormal development. The transcriptional and developmental analysis suggested that Kv2.1 subunits antagonize each other during RF formation (Table 2; Fig. 1).

The member of Li-CAM family member of the same family - the zebrafish Chl1a/Camel has been implicated in Sspo secretion and hydrocephalus (Yang et al., 2021). The Kcnb1 LOF mutation causes an increase of Sspo production (Fig. 1B), similar to the Camel/Chl1a GOF (Yang et al., 2021). Further supporting the link between Kv2.1 and Camel/Chl1a, the level of *camel/chl1a* transcripts increased in the Kcnb1 mutant (Table 2). In contrast, the Camel LOF was reduced or even abolished the RF in line with reduction of *sspo* and *camel* expression caused by the *kcng4b* mutation (Table 2; Fig. 1C, C’). These results illustrate the inverse developmental regulatory link between Kv2.1 and Chl1a.

Previously the electron microscopy data revealed several phases of RF organization in amphibians and mammals with the RF assembled from multiple individual microfilaments (Nualart and Hein, 2001; Pérez-Fígares et al., 2001; Sepúlveda et al., 2021; Woollam and Collins, 1980; Yulis et al., 1998). Here the following steps of reorganization of Sspo into RF were presented: a) the secreted RF material on the SCO’s apical surface, b) the formation of extended filaments, c) the merger of filaments in the CA resulting in the RF formation (Fig. 1). After the RF cut, the “flower-like” ends form (Bellegarda et al., 2023), which may represent the dissociation of individual microfilaments. If this assumption is correct, the RF assembly appears to be reminiscent of a rope assembled from individual threads.

CSF flow has been implicated in RF formation (Lang et al., 2006; Picketts, 2006). Like the subterranean river, the CSF flows through the BVS cavities consisting of the narrows, expanses and straights (Sherman and Citrin, 1986). The neuroanatomical location of the SCO at the anterior end of the CA, i.e. one of the BVS narrows, suggests an increase in flow turbulence at this location (Sepúlveda et al., 2021). The second BVS narrow is at the posterior end of CA. It is associated with the FO. The third is located at the entrance to central canal (*obex*), which is aligned with another CVO, the area postrema (AP) (García-Lecea et al., 2017; Korzh and Kondrychyn, 2020). The areas of mid-CA, and vIV likely represent the areas of low turbulence. The longest stretch of laminar CSF flow corresponds to the relatively strait and uniform in diameter central canal, where laminar CSF flow would be expected (Fig. 3). Thus, at least three CVOs - SCO, FO and AP - could be associated with predicted areas of increased CSF turbulence. The RF is attached to the SCO and FO and this attachment defines the RF trajectory. In particular, the SCO and FO anchor the aRF, while the FO anchors the pRF. As a tense and oscillating string (Bellegarda et al., 2023; Woollam and Collins, 1980), the RF likely to generate the most tension in areas of increased turbulence, which may explain a need for RF attachment identified nearly 50 years ago (Woollam and Collins, 1980).

Changes in microfilament distribution and curvature of the SCO apical surface further illustrate the viscoelastic effects of RF on the SCO morphogenesis previously reported in the zebrafish and rodents (Xu et al., 2023; Yang et al., 2021). Perhaps, this effect could be attributed to the pulling effect of CSF flow along the anterior-posterior (A-P) axis (Jeong et al., 2024). In such a scenario, the RF likely transmits the pull of CSF flow to the SCO which may cause the posterior-ward SCO elongation and dorsal-ward FO elongation (Fig. 2, 3). The RP cells secrete BMP known to increase tissue fluidity (Butler and Dodd, 2003; Shinozuka and Takada, 2021; Yang et al., 2023). The significant mechano-elastic stretch of the RP during primitive lumen reduction has been reported in rodents and zebrafish (Kondrychyn et al., 2013; Korzh, 2014; Ševc et al., 2009; Snow et al., 1990), providing direct evidence for the high fluidity of the RP cells compared to that of FP cells in the spinal cord.

As a derivative of the RP, which undergoes a process of stretched morphogenesis (Kondrychyn et al., 2013; Korzh, 2014), the SCO likely shares significant mechano-elastic fluidity with the roof plate. Notably, the anterior FP, especially the FO, also stretches significantly in contrast to the spinal cord FP. One reason for this may be the expression of the “dorsal” Wnts such as Wnt3 in the anterior FP (Cathleen Teh et al., 2015). In the absence of the aRF caused by the defective Wnt/β-catenin signaling (Fig. 4), the SCO fails to extend posteriorly. A similar phenotype was observed in the Chl1a- or Fabp7b-deficient zebrafish (Jeong et al., 2024; Yang et al., 2021) and Sspo-deficient mouse (Xu et al., 2023). Previously, it has been shown that the absence of RF in several zebrafish mutants leads to the curly-down phenotype, which is reminiscent of scoliosis (Cantaut-Belarif et al., 2018; Grimes et al., 2016; Kramer-Zucker et al., 2005; Lehmann and Naumann, 2005). This finding is in contrast to the results of the local RF ablation at the level of the spinal cord, which did not induce such a phenotype (Bellegarda et al., 2023). This could be due to the preservation of anterior region of the RF, which during development was “threaded” through the BVS “narrows”, so that the continuous elongation of the RF will eventually restore it. The RF “threading” through CA, MHB and *obex* appears to be critical for RF recovery (Lang et al., 2006; Picketts, 2006). The same purpose could be served by the existence of separate regions in the RF limited by their attachment to CVOs - SCO – FO, FO - ampulla terminalis (Fig. 3). Focal RF ablation in the spinal cord between the FO and the ampulla terminalis does not affect the SCO-FO region. Therefore, it may not have the same negative effects as the aRF deficiency.

SCO and FO have very different developmental history. They belong to the counteracting axial signaling centers, i.e., the FP and RP, respectively. Although FO and SCO play different functions during RF development, they share some common features, such as the expression of Sspo and Wnt3 (Clements et al., 2009; Lehmann and Naumann, 2005; López-Avalos et al., 1997; Teh et al., 2015). Importantly, in the context of RF development, the aRF and pRF both attach to the FO and maintain this attachment (Woollam and Collins, 1980) despite the separation of the RF and hindbrain FP caused by the ventral bending of the latter (Fig. 1-3). In this respect the FO differs from the posterior FP. To our knowledge no FO-specific developmental regulators have been found so far. It seems that one of the developmental roles of the FO is to keep the RF trajectory straight between the SCO and the entrance of the central canal. Thus, in addition to the synthesis and secretion of Sspo, FO may play a critical role in the RF assembly and maintenance of mechanical stability.

Since several Wnts and Sspo are co-expressed in the SCO (Fig. 4) (Clements et al., 2009; García-Lecea et al., 2017; Krauss et al., 1992; Molven et al., 1991), the question arose whether the secretion of these proteins is dependent on Kv2.1 activity. Previously, it was shown that the Wnt secretion in the zebrafish brain may involve the cholesterol-enriched regions (Ng et al., 2016). The analysis of Wnt3 secretion shows that the aRP may be the source and target of Wnt3 (Cathleen Teh et al., 2015; Veerapathiran et al., 2020). Thus, the SCO development may require both cell-autonomous and non-cell-autonomous Wnt/β-cat signaling involving Wnt3. Like Wnt3 secretion, Sspo secretion (Fig. 6) is likely to rely on cholesterol-rich regions of the plasma membrane. It has been proposed that the lipid rafts are surrounded and maintained by the clusters of electrically inactive Kv2.1 (Deutsch et al., 2012; Johnson et al., 2019; Tamkun et al., 2007; Xia et al., 2004).

Kv2.1 has been implicated in the negative regulation of secretion at presynaptic compartments (Dodson and Forsythe, 2004) and insulin secretion by pancreatic β-cells (Jacobson et al., 2007). Cholesterol depletion or Kv2.1 knockout causes an increase in glucose-stimulated insulin secretion (Jacobson et al., 2007; Rorsman and Ashcroft, 2018; Xia et al., 2004). The mechanisms of insulin and Sspo secretion appear to share other common features. Heterozygous variants in the hepatocyte nuclear factor 1a (HNF1a) cause MODY3 (maturity-onset diabetes of the young, type 3 by inducing ER stress in pancreatic β-cells and decreasing their number, insulin expression, and secretion) and Dandy-Walker malformations, including hydrocephalus (Chen et al., 2022; Matsukura et al., 2017). This suggests that HNF1a may be a common element in the developmental mechanisms of BVS and pancreatic β-cells. Notably, the panembryonic RNAseq has shown that the expression of *hnf1a* and *insb* is significantly reduced in the Kcng4b-deficient zebrafish (Jędrychowska et al., 2024) suggesting that HNF1a may be involved in the Kv2.1-mediated processes of insulin secretion by the pancreas and Sspo secretion by the SCO.

Our study demonstrated the role of the voltage-gated potassium channel Kv2.1 and its subunits (Kcnb1 and Kcng4b) in the formation of the RF. The latter may be one of the reasons for the quantitative changes in the BVS of zebrafish lacking Kv2.1 subunits. Kv2.1 is essential for the regulation of Sspo secretion, with cholesterol playing a key role in this process. RF tension drives the morphogenesis of SCO and FO.

## Materials and Methods

### Animals

Zebrafish (*Danio rerio*) were maintained according to established protocols (Westerfield, 2007) in the Zebrafish Core Facility at the International Institute of Molecular and Cell Biology in Warsaw (licensed breeding and research facility, PL14656251, registry of the District Veterinary Inspectorate in Warsaw; 064 and 051: registry of the Ministry of Science and Higher Education). All the experiments with zebrafish embryos, larvae and adults were performed in accordance with the European Communities Council Directive (63/2010/EEC). Developmental stages in hours post-fertilisation (hpf) are based on (Kimmel et al., 1995). *kcnb1sq301/sq301* (also referred to as kcnb1^-/-^) mutant was described previously (Shen et al., 2016). *kcng4b-c1^-/-^* mutant (waw304) was described previously (Gasanov et al., 2021).

The transgenic lines were used in this study as the *in vivo* markers: ET33-mi2A transgenic line (sqet33mi2AEt) with a transposon insertion in the position of *prom1a* (García-Lecea et al., 2017), expresses cytosolic green fluorescent protein (GFP) in the roof plate, SCO, ear, etc.; Wnt3-GFP (sq19Tg) expresses cytosolic GFP in the roof plate, SCO, etc. (Cathleen Teh et al., 2015), KR15-16 (sqKR15ct16Tg) expresses membrane-bound KillerRed at the midbrain-hindbrain boundary in the En1b-like manner (Teh et al., 2010)).

### Microscopy

#### Live imaging

Zebrafish embryos were raised in E3 medium (2.5 mM NaCl, 0.1 mM KCl, 0.16 mM CaCl2, and 0.43 mM MgCl2) with the addition of 0.2 mM 1-phenyl-2-thiourea (PTU, Merck, Germany) to block pigmentation. At selected developmental stages, embryos were manually dechorionated, anesthetized with 0.02% tricaine (Sigma-Aldrich, USA) and oriented by embedding in 2 % methylcellulose (Merck, Germany) on the glass slides. A research stereomicroscope SMZ25 (Nikon, Japan) was used to image acquisition.

### Light-sheet confocal microscopy (LSCM) imaging

In vivo imaging and imaging of fixed specimens were performed as previously described (Jedrychowska et al., 2021). For imaging of fixed material, 0.8 % low-melting agarose in phosphate buffer saline (PBS) was used instead of E3 0.02 % tricaine medium. Zeiss Lightsheet Z. 1 with W Plan-Apochromat 20x/1.0 UV-VIS (for in vivo imaging), 40x/1.0 UV-VIS or 63x/1.0 UV-VIS (for fixed embryos) objectives were used, transmitted LED light was also used to obtain high-resolution bright-field images of zebrafish ear. Data were saved in the LSM or CZI format and processed using ZEN (Zeiss) or ImageJ 1.51n (Fiji) software. Maximum intensity or sum slices projections were generated for each z-stack. Brightness/contrast adjustments and resizing were performed using FastStone viewer 7.4 (FastStone Soft).

### Immunohistochemistry

The embryos were stained by the two-color immunohistochemistry for RF using polyclonal rabbit AFRU antibody (1:1000) (a gift of Drs. J. Grondona [Malaga, Spain], E. Rodriguez, and M. Guerra[Valdivia, Chile]) according to the described protocol (Korzh et al. 1998). The GFP detected was expressed in transgenic embryos.

### qRT-PCR

The SsoAdvanced Universal SYBR Green Supermix and CFX Connect real-time PCR system (Bio-Rad, USA) was used for the Quantitative Real-Time Polymerase Chain Reaction (qRT-PCR) according to the manufacturer’s instructions. The TRIzol-chloroform RNA extraction protocol (Sigma-Aldrich, USA) was used to extract total RNA from 20 to 50 zebrafish embryos at 28, 48, and 72 hpf. RNA concentration was assessed using a NanoDrop™ 2000 spectrophotometer (Thermo Scientific, USA), and cDNA was synthesized from 1 μg of RNA using the iScript™ Reverse Transcription kit (Bio-Rad, USA). qRT-PCR was performed using gene-specific primers for eef1a1l1, the reference reaction for the housekeeping gene, and other primers (Suppl. Table 1) were used to amplify mRNAs of interest selected based on results of the RNAseq analysis described above. The threshold cycle of each target and reference gene amplification in control and mutant embryos was determined automatically by LightCycler® 96 Software (Roche Diagnostics, USA). Fold change in mutants versus controls was calculated with delta-delta-C(t) method and Student’s two tailed t-test with respect to the mismatch control was used to determine statistical significance. Statistical analysis including standard deviation calculation, was performed using Microsoft Excel (Microsoft, USA) and GraphPad Prism 5 (GraphPad, USA) software. Melting curve analysis and agarose gel electrophoresis were performed as product specificity controls, while the reaction efficiency (E) was calculated separately for each gene analyzed.

### Cholesterol depletion

Zebrafish embryos were raised in Petri dishes containing embryo media and maintained at 28°C. Prior to treatments, partial tearing of the chorions was performed to facilitate exposure to a chemical. A total of 20 embryos were transferred to 25 mL of embryo media in glass Petri dishes.

For cholesterol depletion, embryos were exposed to methyl-β-cyclodextrin (mβCD) at 48 hours post-fertilization (hpf), prepared to a final concentration of 2.5 μM. The treatment was applied for 40 minutes, after which embryos were washed three times with cold phosphate-buffered saline (PBS) for 5 minutes each. Following the washes, embryos were fixed in 4% paraformaldehyde (PFA) for further analysis.

To block cholesterol biosynthesis, Atorvastatin was applied at a final concentration of 10 µM. Treatment was initiated at 24 hpf, and 20 embryos were incubated in the solution for 24 hours. After the treatment, embryos were washed and fixed, following the same procedure as described for the mβCD treatment.

The control groups were maintained in embryo media (E3 media) without any treatment, and for the mβCD treatment, an equivalent amount of DMSO was used as the vehicle control. All experimental conditions, including both mβCD and atorvastatin treatments, were repeated three times to ensure reproducibility.

### In Vivo Optogenetics for Light-Induced Oxidative Stress

The 24 hpf embryos of KR15-16 transgenic line were mounted in 1% low-melting point agarose and exposed to maximum intensity of green light (UV filter set 38HE, 68 mW, 1x/3.2 objective from the Nikon fluorescent upright SMZ25 microscope for 90 min according to a published protocol (Teh et al., 2010; Teh and Korzh, 2014). The 48 hpf embryos were processed for AFRU IHC and LSCM.

### AFRU antibody injection

2 nl of the whole AFRU antibody (in contrast to the partially purified and 2 x concentrated antibody used by (Rodríguez et al., 1990) was injected into the IVth (hindbrain) brain ventricle of 24 hpf embryos (Shen et al., 2016)). The 48 hpf embryos were processed for AFRU IHC and LSCM.

## CRediT authorship contribution statement

Razieh Amini: Methodology, Investigation. Ruchi Jain: Methodology, Investigation. Vladimir Korzh: Writing – review & editing, Writing – original draft, Validation, Supervision, Project administration, Methodology, Investigation, Funding acquisition, Data curation, Conceptualization.

## Declaration of generative AI and AI-assisted technologies in the writing process

During the preparation of this work the author(s) used DeepL Write to improve language and readability. After using this tool/ service, the authors reviewed and edited the content as needed and took full responsibility for the content of the publication.

## Data availability

Data will be made available on request.

## Acknowledgements

The authors thank Prof. Jacek Kuznicki and all members of the Laboratory of Neurodegeneration, the Microscopy and Zebrafish Core Facilities of the International Institute of Molecular and Cell Biology in Warsaw. The draft of this paper has been deposited to BioRxiv, XXX.

VK acknowledges support from the Opus grants of the National Science Centre (NCN), Poland (2020/39/B/NZ3/02729).

## Supplementary Figures

Fig. S1. The RF connects the SCO, FO and projects towards the central canal in a straight line. A - Tg (Sspo-GFP), the white arrows show the weak GFP signal at the hindbrain FP apical surface; B - ET33-10 (modified from Garcia-Lecea et al., 2017; green - GFP; red - projected RF trajectory). Arrowhead – GFP-Sspo present ventrad to the pRF.

Fig. S2. aRF development could be blocked by ROS or AFRU antibody.

During development KillerRed (KR) is expressed at the MHB (asterisk) of Tg(KR15-16) transgenics in the En1b-like manner (A, B; Irradiation with intense green light (24 hpf) triggered production of ROS, which blocked aRF formation (C, D). A similar outcome has been caused in some embryos after injection of the AFRU antibody (E, F), which was previously shown to have the neutralizing activity in mammals (Rodriguez et al., 1990). (+) - increased background staining probably due to remnants of injected AFRU antibody. Scale bar - 20 um.

