## Supplementary figures and images for "Kcnb1-Kcng4 axis regulates Scospondin secretion and Reissner fiber development"

### Supplementary Figures 1-2

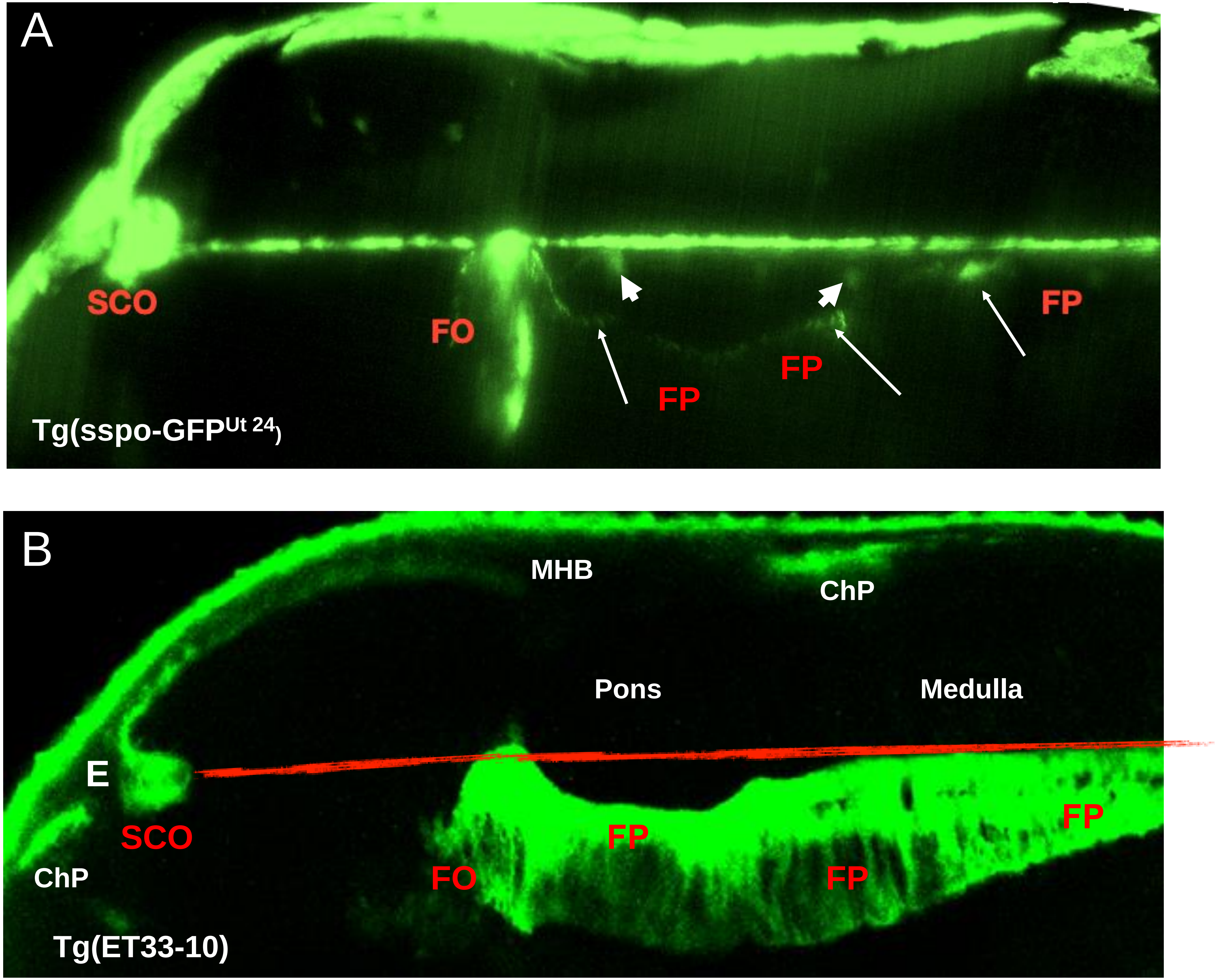

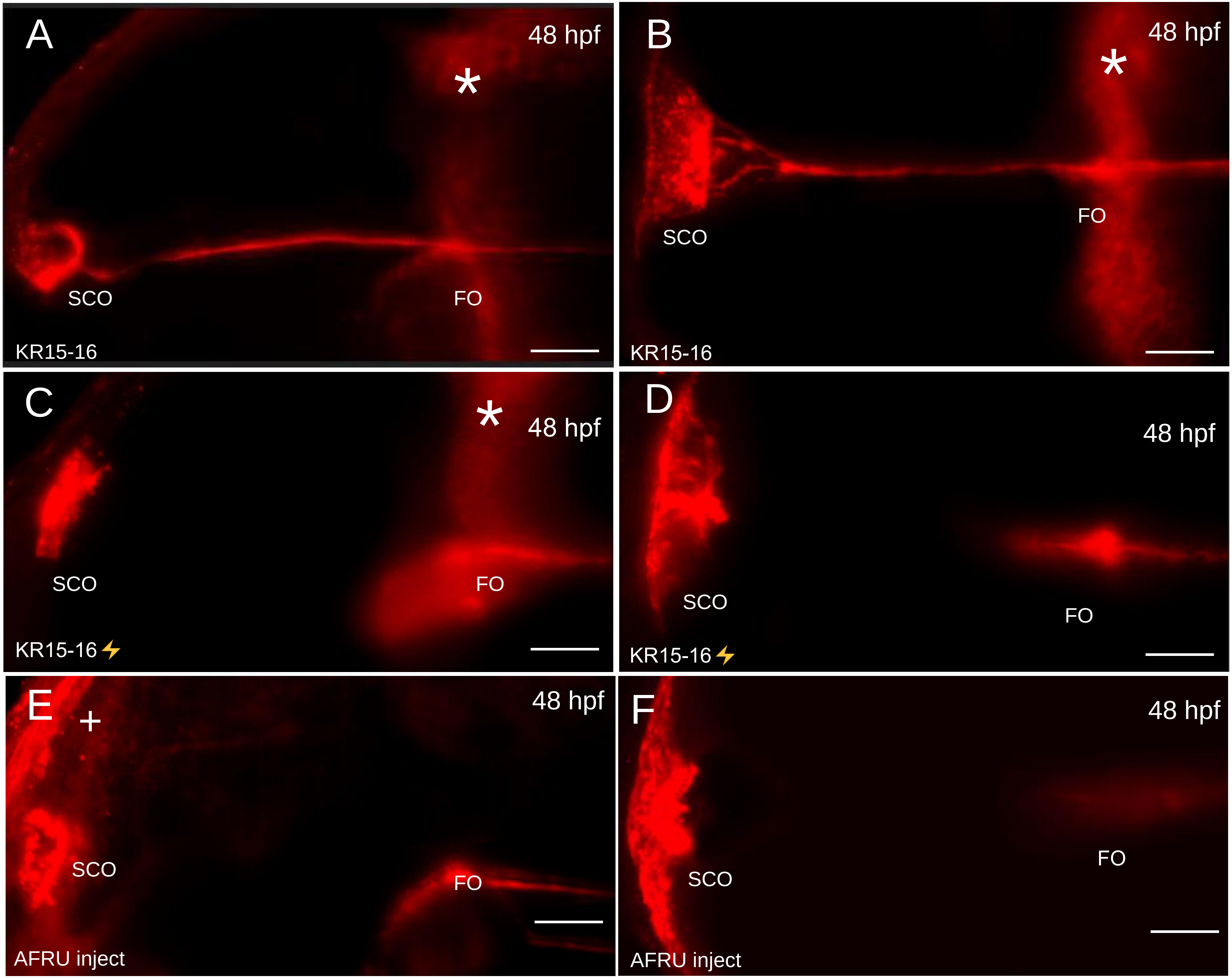
